# Developmental diversification of cortical inhibitory interneurons

**DOI:** 10.1101/105312

**Authors:** Christian Mayer, Christoph Hafemeister, Rachel C. Bandler, Robert Machold, Kathryn Allaway, Xavier Jaglin, Renata Batista Brito, Andrew Butler, Gord Fishell, Rahul Satija

## Abstract

Diverse subsets of cortical interneurons play a particularly important role in the stability of the neural circuits underlying cognitive and higher order brain functions, yet our understanding of how this diversity is generated is far from complete. We applied massively parallel single-cell RNA-seq to profile a developmental time course of interneuron development, measuring the transcriptomes of over 60,000 progenitors during their maturation in the ganglionic eminences and embryonic migration into the cortex. While diversity within mitotic progenitors is largely driven by cell cycle and differentiation state, we observed sparse eminence-specific transcription factor expression, which seeds the emergence of later cell diversity. Upon becoming postmitotic, cells from all eminences pass through one of three precursor states, one of which represents a cortical interneuron ground state. By integrating datasets across developmental timepoints, we identified transcriptomic heterogeneity in interneuron precursors representing the emergence of four cardinal classes (Pvalb, Sst, Id2 and Vip), which further separate into subtypes at different timepoints during development. Our analysis revealed that the ASD-associated transcription factor Mef2c discriminates early Pvalb-precursors in E13.5 cells, and removal of *Mef2c* confirms its essential role for Pvalb interneuron development. These findings shed new light on the molecular diversification of early inhibitory precursors, and suggest gene modules that may link developmental specification with the etiology of neuropsychiatric disorders.

## INTRODUCTION

Cortical interneurons (cINs) are an extremely diverse group of inhibitory cells that vary widely in morphology, physiology, connectivity, and molecular markers^1–5^. In the visual cortex alone, recent studies have identified in excess of 20 transcriptionally defined interneuron clusters, which represent specific subtypes or states^6,7^. A majority of cINs are derived from progenitor cells residing in transient developmental ventral brain structures known as the medial and caudal ganglionic eminences (MGE and CGE, respectively). Progenitors from these GEs develop into distinct neuronal subtypes that produce non-overlapping populations of cINs (i.e. MGE-type (Pvalb and Sst-expressing) vs. CGE-type (Vip and Id2/Ndnf-expressing) cINS)^8^. However, the molecular heterogeneity across cells within these same eminences remains less well characterized, particularly as most neurochemical markers that define cIN subtypes are not expressed at early timepoints^9^. While known transcription factors (TFs) show heterogeneous expression patterns within immature cINs^10–12^, unsupervised analyses of cellular heterogeneity have the potential to shed new light on early fate transitions and decisions^13,14,15^ driving the establishment of cIN subtypes, and to improve our understanding of how the vast diversity in adult cINs emerges from a small pool of mitotic progenitors^16–18^. Here, we combine single cell RNA-sequencing approaches (scRNA-seq; plate-based RNA-seq; Drop-seq^19^, 10x Genomics platform) with genetic fate mapping techniques. Our analysis revealed three fundamental insights. First, diversity within mitotic progenitors is largely accounted for by cell cycle state and differentiation that are seeded with eminence specific transcription factors. Second, soon after cells become postmitotic three pan-eminence transcriptional programs (representing one interneuron and two projection neuron precursor states) emerge, and in parallel, eminence-specific transcriptional programs escalate. Third, by integrating datasets across a developmental time course, we show how the embryonic interneuron precursor state evolves into the diversity seen in the adult. This analysis has enabled us to connect the transcriptional states of these cells from their inception to their mature adult identities, and to identify and validate new early markers of subtype specification.

## RESULTS

### Transcriptional profiling of GE cells

We manually dissected the MGE, CGE and LGE of E13.5 (MGE) and E14.5 (CGE, LGE) wild type mouse embryos, embryonic stages corresponding to periods of abundant neurogenesis within each of these structures^20–23^. We collected the ventricular zone (VZ), subventricular zone (SVZ) and adjacent mantle zone (MZ) within each eminence (Fig. 1A), which includes both symmetrically and asymmetrically dividing mitotic progenitors, as well as postmitotic precursor cells.

**Figure 1.**
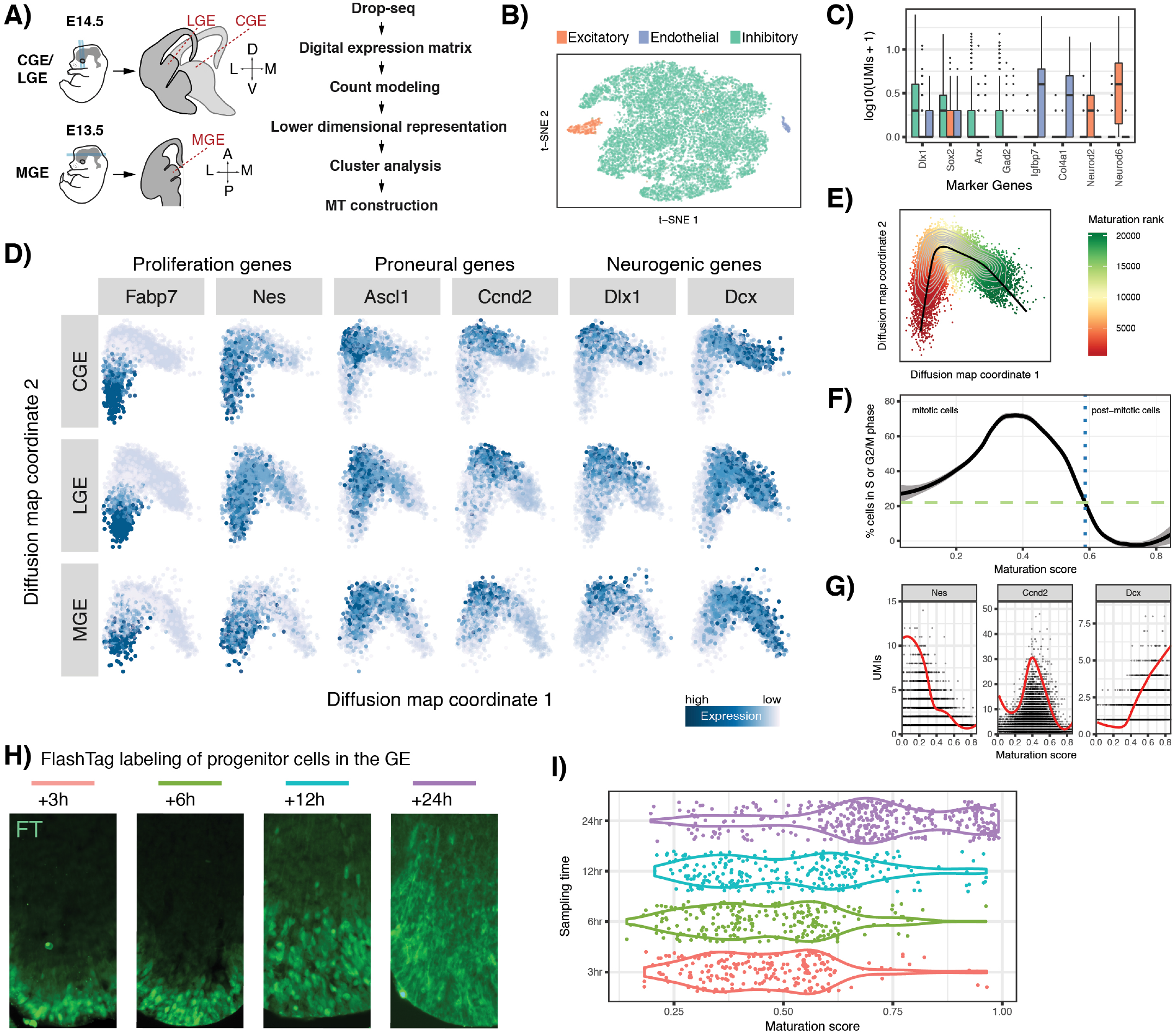
Transcriptional heterogeneity in the GEs. **A)**Schematic of experimental workflow. MGEs were dissected from horizontal brain sections, whereas CGEs and LGEs were dissected from coronal brain sections, as indicated by the light blue lines. Axes: Dorsal (D), Ventral (V), Posterior (P), Anterior (A), Lateral (L), Medial (M). **B)** Visualization of Drop-seq data using t-SNE, depicting the presence of distinct minority cell types. **C)** Expression of canonical marker genes shows the vast majority of cells identified were GE precursors (green), while a small minority of cells represented excitatory neurons (orange) and vascular endothelial cells (blue) based on canonical marker expression. **D)** Diffusion map analysis of eminence datasets suggests a developmental continuum. Each eminence was analyzed independently, revealing nearly identical patterns. Cells are colored according to the expression of canonical regulators. **E)** A principal curve was fitted to the two dominant diffusion map coordinates to order cells along a maturation trajectory (MT). **F)** The percentage of actively cycling cells transitions during maturation (dotted-blue line), marking the shift during MT into a postmitotic phase. **G)** Expression (molecules/cell) of canonical regulators, as a function of the position along the maturation trajectory. Curve reflects local averaging of single cell expression. **H)** Coronal brain sections of the ganglionic eminences, as cells migrate away from the VZ (Ventricular Zone). Images were taken 3, 6, 12 and 24 hours after fluorescent labeling with FlashTag technology. **I)** Maturation score distributions from the FlashTag dataset, separated by timepoint post labeling (3 (red), 6 (green), 12 (blue) and 24 (purple) hours). Cells from later time points were positioned further along the inferred maturation trajectory.

After cell dissociation, we utilized Drop-seq^19^ to sequence the transcriptomes of 5,622 single cells from the MGE, 7,401 from the CGE, and 8,543 from the LGE, using three independent biological replicates for each eminence. We sequenced libraries to an average depth of 15,990 reads/cell, detecting on average 1,626 UMIs, comparing favorably to a recently published Drop-seq dataset focused on retinal bipolar cell heterogeneity^24^. We first regressed out confounding sources of technical variation between single cells, including sequencing depth and library complexity (Supplementary Methods). We then used a curated gene set^25^ to define a cell-cycle score for each cell in the dataset, allowing the assignment of a mitotic (M, S, G-phase) or postmitotic status (cell cycle score near zero) to every cell (Supplementary Fig. 1). This enabled us to perform latent variable regression to mitigate heterogeneity resulting from cell-cycle state^26^, so that subsequent analyses would not be dominated by mitotic phase-specific gene expression.

To identify distinct groups of cells in our data, we applied a graph-based clustering algorithm after non-linear dimensional reduction with diffusion maps^27^. We identified small populations of excitatory neurons *(Neurod6;* 2.6% of cells) and endothelial cells (*Igfbp7*; 0.7% of cells) (Fig. 1B, C), both of which were excluded from further analysis. The remaining 96.7% of cells were GE-derived progenitors and precursors (e.g., *Dlx1*; Fig. 1B, C). Within this population, the expression of early, intermediate, and late marker genes was strongly associated with the top diffusion map coordinates (Fig. 1D). The same result was obtained using principal component analysis (Supplementary Fig. 1).

To establish a quantitative temporal account of differentiation programs within each eminence, we implemented an unsupervised procedure to ‘order’ single-cells based on their expression profiles. After calculating the diffusion map coordinates (DMC), we fit a principle curve through the resulting point cloud which summarizes the optimal path through a continuous landscape of differentiation^28^. We hypothesized that projecting cells onto this curve represents progression along a developmental trajectory (Fig. 1E). We confirmed that our resulting maturation trajectory (MT) was reproducible with a complementary technique to calculate ‘pseudotime’ (Supplementary Fig. 1)^13^, alongside multiple validations. First, based on their previously calculated cell-cycle scores, we performed a supervised analysis and observed that cells early in the MT were mitotic, whereas late cells were postmitotic (Fig. 1F). Second, gene expression dynamics of canonical regulators along the MT strongly recapitulated known dynamics associated with neuronal maturation (Fig. 1D, G)^29,30^. Finally, we utilized FlashTag technology^31^ to fluorescently label cells in the VZ^32^ of the GEs, and performed scRNA-seq on cohorts of 3, 6, 12 and 24 hour-old neurons as they migrated away from the ventricle (Fig. 1H). We found that neurons born at these sequential timepoints were similarly distributed progressively along the MT timeline, thereby independently confirming the MT with real time (Fig. 1I, Supplementary Fig. 2).

MT enabled us to specifically isolate mitotic progenitors and compare their gene expression across the three GEs. Consistent with previous studies, we found a small number of transcription factors enriched in particular eminences (Fig. 2A, Supplementary Fig 3, 4, Supplementary Table 1), many of which have been previously described^10^. The majority of dynamically expressed genes followed a similar temporal sequence across all eminences with three sequential waves of gene expression (Fig. 2B,C, Supplementary Table 2), consistent with findings in excitatory progenitors of the cortex^32^. In-situ hybridization (ISH) confirmed that these waves describe the sequential expression of stem-cell (e.g. *Nes*), transient amplifying (e.g., *Ccnd2*), and postmitotic genes (e.g. *Dcx*; Fig. 2D)^33^, roughly correlating with the spatiotemporal progression from the VZ to the MZ (Supplementary Fig. 5). To further quantify these results, we calculated the primary sources of transcriptional variance in these progenitors (Supplementary Methods), and found that developmental progression and cell cycle were the dominant sources of variance, with maturation proportionally explaining six-fold more variance compared to eminence-of-origin (Fig. 2E).

**Figure 2.**
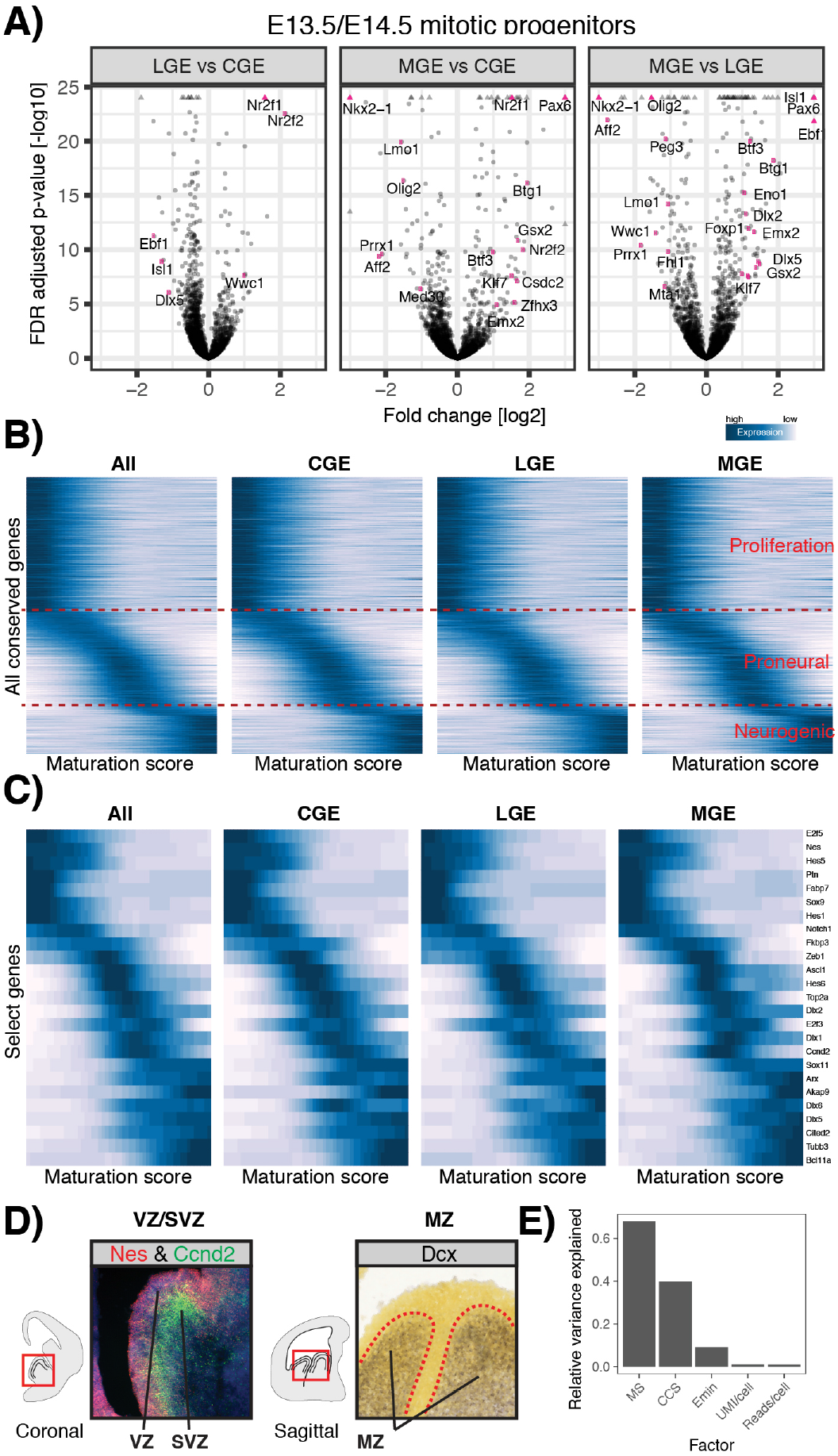
A common developmental program of gene expression functions in mitotic progenitors of all three GEs. **A)** Volcano plots depicting differential gene expression across eminences for early mitotic cells (MT < 0.3). Gene names of transcription factors are annotated. **B+C)** Gene expression dynamics in mitotic cells, based on local averaging of single cell data, plotted along MT for all (B) and select (C) developmentally regulated genes. These dynamics reveal sequential waves of gene expression that are consistent between eminences. **D)** In-situ hybridization (ISH) patterns of early, intermediate and late MT genes in the GEs that are highly expressed within anatomical boundaries of the Ventricular Zone (VZ), Subventricular Zone (SVZ), and Mantle Zone (MZ), respectively; The ISH images for Dcx was taken from the Allen Brain Institute. **E)** We calculated the relative variance explained individually by a set of annotated factors (maturation score (MS), cell cycle score (CCS), eminence of origin (Emin), unique molecular identifiers per cell (UMI/cell), and reads per cell (reads/cell)) and normalized this to the variance explained by the first principle component (Supplementary Methods). Developmental progression and cell cycle stage (CCS) were the dominant sources of variance in mitotic cells, with maturation relatively explaining six-fold more variance compared to eminence-of-origin. Technical factors (UMI/cell and reads/cell) contributed minimal relative variance.

We next explored whether transcriptional programs drive the emergence of distinct progenitor or precursor cell types in the GEs. To detect potential fate divergence of cells along the MT, we bootstrapped the construction of a minimum spanning tree (MST)^34^ (Fig. 3A; Supplementary Methods), and summarized the combined result using multi-dimensional scaling. We found that as cells become postmitotic, the MST from each GE divides into three branches (Fig. 3B; Supplementary Methods). Notably, despite the presence of inter- and intra-GE specific gene expression (Fig. 2A), we did not observe transcriptomic evidence of similar bifurcations in mitotic cells (Supplementary Fig. 6A). Specifically, even supervised analysis using known branch markers, or by profiling 400 mitotically enriched MGE cells sequenced with a modified version of the Smart-seq2 protocol^35^ and sequenced at significantly higher depth (average of 4,130 genes/cell; Supplementary Fig. 6B, C) failed to provide evidence for branching within mitotic GE progenitors. Thus, while a small number of regionally-restricted transcription factors are differentially enriched in distinct populations of mitotic progenitors, transcriptomic branching occurs concurrently with the transition from mitotic progenitors into postmitotic precursor cells.

**Figure 3.**
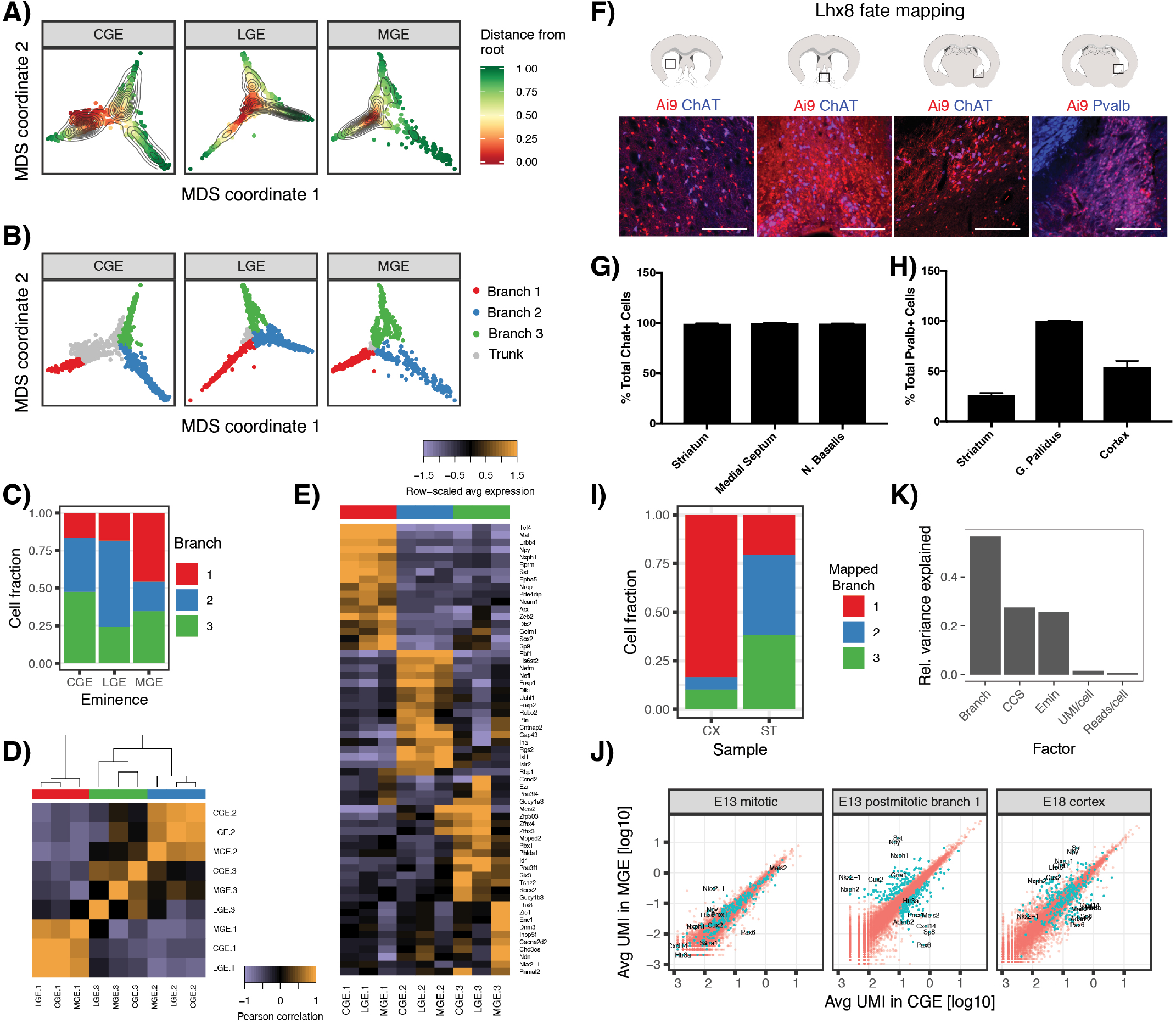
Postmitotic cells from all eminences pass through one of three precursor states. **A)** Multidimensional scaling (MDS) based on the consensus MST depicts a common trunk and three divergent developmental trajectories along MT. Branching trajectories were calculated for each eminence individually. **B)** MST traversal assigned cells to the major branches; trunk, grey; branch 1, red; branch 2, blue; branch 3, green. **C)** While each eminence exhibits three developmental branches, each has varying quantitative contributions to each segment. **D)** Heatmap of the Pearson Correlation of branch averages. Gene expression was highly correlated across eminences for branches 1 and 2. MGE cells for branch 3 exhibited lower correlation with CGE/LGE cells. **E)** Heatmap depicting the top transcriptomic markers for each branch; expression for all cells with the same eminence and branch annotation were averaged together. **F)** Colocalization of Lhxδ-Cre/cerulean; Ai9 (tdtomato) with choline acetyltransferase (ChAT) in the striatum, medial septum, and nucleus basalis, and with Pvalb in the globus pallidus (from left to right). Scale bars = 300 μm. **G)** The percentage of total ChAT÷ cells in the striatum, medial septum, and nucleus basalis that are labeled with tdtomato in Lhx8-Cre/cerulean; Ai9 mice. n=2 mice, 8 sections per brain. Error bars indicate standard deviation across brains. **H)** The percentage of total Pvalb-positive cells in the striatum, globus pallidus, and cortex that are labeled with tdtomato in Lhx8-Cre/cerulean; Ai9 mice. n=2 mice, 6 sections per brain. Error bars indicate standard deviation across brains. **I)** Mapping of E18.5 cortical (CX) and subcortical (ST) cells to each of the three E13.5/E14.5 branches using marker gene expression. Barplot shows that cortical E18.5 cells map preferentially to branch **1**, consistent with its annotation as an interneuron precursor state. Supplementary Figure 9D-E shows genes conserved between E18.5 cells and E13.5/E14.5 branches. **J)** Differential expression analysis between MGE and CGE postmitotic cells in the interneuron precursor stage. Scatter plot (middle) denotes average expression of all genes, with differentially expressed genes in blue. These genes tend to continue to be differentially expressed between MGE and CGE-derived populations at E18.5 (right), but are largely expressed at equal levels across eminences in E13.5 mitotic progenitors (left). **K)** Relative variance calculation as in Fig. 2e, for postmitotic cells (Supplementary Methods). Branching annotations are the dominant sources of variance, with eminence-of-origin playing an increasing role as well. Residual cell cycle variation is due to our conservative cutoff for the mitotic/postmitotic transition (Figure 1F).

We assigned cells to branches by traversing the final MST and annotating major splits (Fig. 3B, C). Strikingly, even though branched trajectories for each eminence were calculated independently, gene expression of branches were highly correlated across eminences (Fig. 3D, E). This finding indicates that, although each GE generates different cell populations, upon becoming postmitotic cells from all eminences pass through one of three conserved precursor states. One group of highly correlated branches (i.e. precursor state 1) expressed known regulators of interneuron development (Fig. 3E), whereas a second group of branches (i.e. precursor state 2) expressed known projection neuron marker genes (Fig. 3E; Supplementary Table 3). The third group of branches (i.e. precursor state 3) exhibited weaker correlation across eminences compared to the precursor states 1 and 2, while appearing to predominantly represent a second projection neuron branch. In particular, the MGE branch differed in its expression of unique marker genes, most significantly in its expression of the transcription factor Lhx8 (Fig. 3E). We genetically ‘fate-mapped’ these cells using a Lhx8-Cre/cerulean driver line and discovered that Lhx8 fate-mapped neurons account for the majority, if not all of the cholinergic projection and interneuron populations within the nucleus basalis, medial septum, and striatum, respectively, as well as the majority of Pvalb-positive projection neurons in the globus pallidus (Fig. 3F-H Supplementary Fig. 6D).

### How does the existence of common precursor states fit into the larger framework of cell-type diversification?

We chose to focus on precursor state 1 to explore the principle of how early precursor states contribute to cell-type diversification. We utilized genetic fate mapping strategies in combination with the 10X Genomics Chromium system for scRNAseq to study cortical interneuron development at embryonic (E13.5, E18.5), postnatal (P10) and adult (P56) stages. Specifically, we used the Lhx6-GFP transgenic mouse line to select for postmitotic MGE cells and to precisely discriminate MGE versus CGE precursor cells at E13.5 (Supplementary Fig. 7). While our 10X dataset at E13.5 focused on postmitotic cells, and did not contain the full developmental continuum cells for trajectory reconstruction, these cells were readily assigned to the three branches identified by Drop-seq, with nearly identical marker expression (Supplementary Fig. 7, Supplementary Methods).

For later stages, when cells have migrated out of the GEs, we used a Dlx6a-Cre; RCE^loxP^ pan-GE fate mapping strategy to collect cortical interneurons at E18.5 and P10 (Supplementary Fig. 8) and utilized the publicly available Allen Brain Institute scRNA-seq dataset^6^ for the adult timepoint. To confirm that cells passing through precursor state 1 give rise to cortical interneurons, we mapped each E18.5 cell to the E13.5/E14.5 precursor states using a correlation-based distance metric (Supplementary Methods). As expected, the vast majority of subcortical cells mapped to projection neuron precursor states (Fig. 3I). By contrast, more than 80% of Dlx6a-positive cortical cells at E18.5 mapped to precursor state 1, based on their expression of canonical regulators of interneuron development (Dlx2; Arx; Maf) (Fig. 3I, J, Supplementary Fig. 9). Of the remaining Dlx6a-positive cortical population, which mapped to precursor states 2 and 3 (Fig. 3I, Supplementary Fig. 9), the majority of neurons were identified as a Meis2-expressing CGE-derived GABAergic population. These cells have been recently described as being present in the cortical white matter^36^ and likely represent projection neurons (Supplementary Fig. 9).

To explore how heterogeneity arose from precursor state 1, we isolated postmitotic cells from this state and performed differential expression analysis on cells originating from the MGE versus CGE. We found 102 genes that were differentially expressed between cells originating from the MGE and CGE (Fig. 3J). Notably, these genes tended to remain differentially expressed in the cortex at later timepoints, but only a small fraction was differentially expressed in mitotic progenitor cells (Fig. 3J). Thus, our data reveal how postmitotic pan-eminence transcriptional programs (precursor states) emerge, and in parallel, eminence-specific transcriptional programs escalate. Consistent with these results, branching trajectories represented the most significant source of variation in these cells, with an increasing contribution from eminence compared to mitotic progenitors (Fig. 3K).

We next asked when adult subtype-specific gene expression patterns first appear during interneuron development. As an adult reference, we applied a graph-based clustering on a publicly available dataset of 3,432 GABAergic P56 cells^6^ (Fig. 4A; Supplementary Methods). We identified 14 prominent inhibitory subpopulations that correspond to known anatomically and physiologically defined subtypes^37,38^ (Fig. 4A, Supplementary Fig. 10). Most of them can be allocated to four major non-overlapping classes of cortical interneurons (Pvalb, Sst, Vip, Id2). In order to link adult heterogeneity to heterogeneity observed at developmental timepoints, we applied our recently developed tool for the integration of scRNA-seq datasets^39^, which aligns shared cell states based on common sources of variation across datasets Fig. 4b-d). We reasoned that if common gene modules were heterogeneous in both adult and developing cells, we may be able to identify early separation of precursors into distinct fates. Indeed, upon integrating P10 and P56 datasets, we observed that P10 cells exhibited strong evidence of transcriptomic separation beyond the four major classes (Fig. 4B). Specifically, they exhibited further subdivisions, including clear segregation within Martinotti and non-Martinotti (X94)^40^ interneurons, as well as Vip bipolar versus multipolar subclasses. Moreover, this revealed clear transcriptomic markers shared between P10 and P56 cells (Fig. 4E; Supplementary Fig. 11-13).

**Figure 4.**
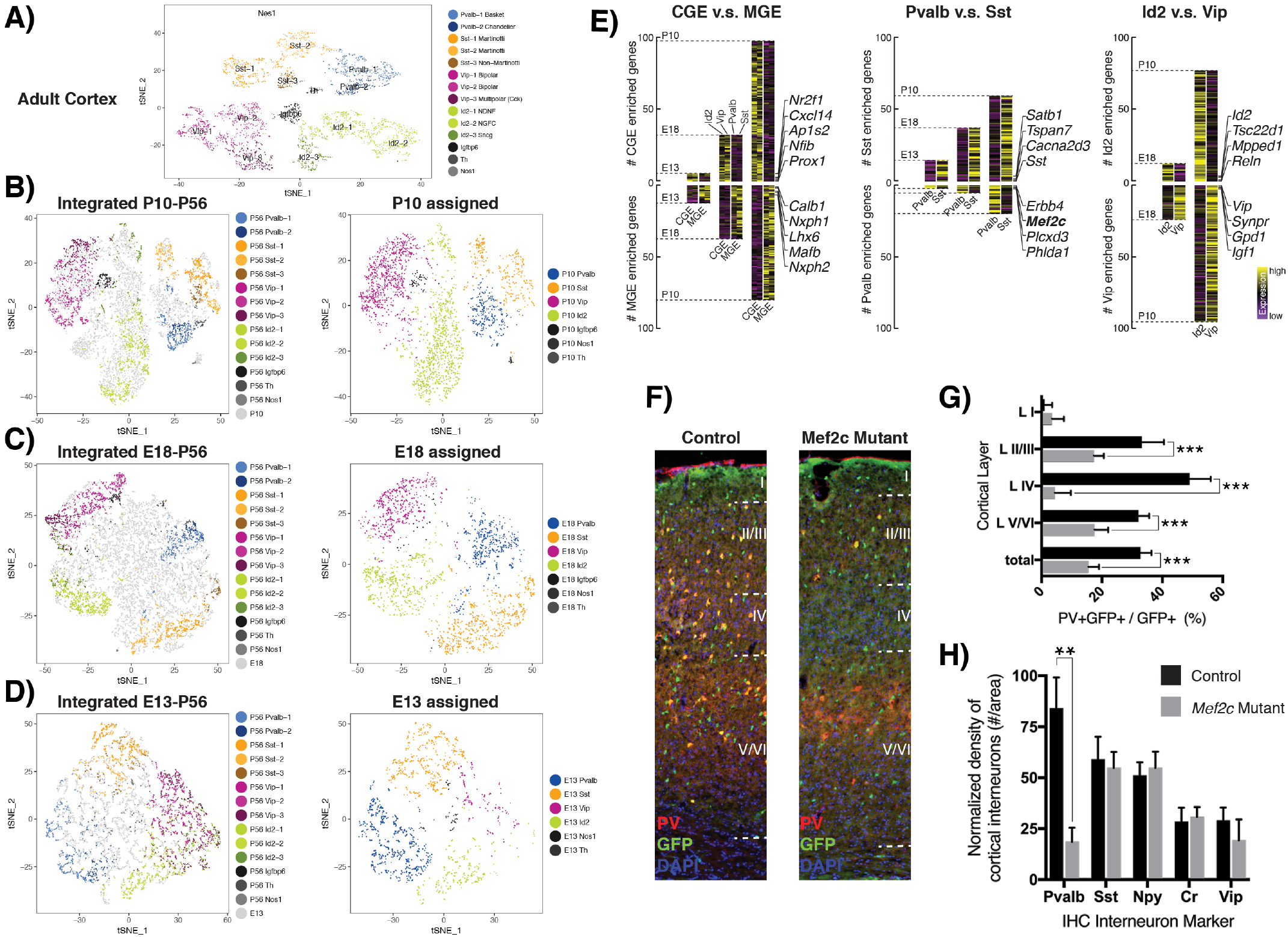
Integrating developmental scRNA-seq datasets to link embryonic heterogeneity to adult interneuron subtypes. **A)** Graph-based clustering of 3,432 interneurons from the adult mouse visual cortex (Allen Institute), reveals 14 clusters, falling into four major classes (Vip/ Id2/ Sst/ Pvalb) with additional subdivisions. Top markers for each cluster are shown in Supplementary Fig. 10. **B)** Integration of 5,176 Dlx6a-Cre;RCE^eGFP^ positive P10 cortical interneurons with the P56 dataset, based on shared sources of variation. P10 cells could be easily assigned to P56 subtypes based on integrated analysis. Conserved markers between P10 precursors and P56 subtypes are shown in (E), with additional subtype-specific markers in Supplementary Fig. 12. (Left) tSNE visualization of all cells, with P56 cells colored by subtype and P10 cells shown in grey. (Right) Same visualization but showing only P10 cells, colored by their subtype mapping (Supplementary Methods). **C)** Integration of 8,382 E18.5 cortical interneurons with the P56 dataset. E18.5 cells separate into the four major classes, with Sst, Vip, and Id2 precursors further separating into subtypes. Additional subtype-specific markers are shown in Supplementary Fig. 13. **D)** Integration of 4,101 interneuron precursor state E13.5 MGE and E14.5 CGE cells with the P56 dataset. E13.5 MGE cells separate into Pvalb and Sst groups while E13.5 CGE cells show only preliminary evidence of initial specification. **E)** Emergence of transcriptomically defined subtypes across development. (Left) Differentially expressed (DE) genes between MGE and CGE derived subsets, that are conserved in both developmental and P56 cells. (Left) DE genes between MGE (which encompasses Pvalb and Sst subsets) and CGE (encompassing Vip and Id2 subsets) classes. Each conserved gene is placed on the heatmap when it is first observed to be DE during development. (Middle, right) same analysis for Pvalb vs. Sst subsets, and Vip vs. Id2 subsets. Earliest conserved genes for each class are annotated, all genes listed in Supplementary Fig. 11. **F)** Conditional deletion of Mef2c in inhibitory neurons using Dlx5/6::Cre;Mef2c^loxp/loxp^RCE and Lhx6i-Cre:Mef2c^loxp/loxp^RCE led to a specific loss of Pvalb interneurons by P20 in cortical layers 2-6. Immunostaining of the P20-P22 somatosensory cortex of Control (left) and Mef2c cKO (right) brains using anti-GFP (green) and anti-Pvalb (red) (DAPI counterstaining shows cortical layers). **G)** Quantification of Pvalb-positive cIN (ratio of the number of Pvalb+ expressing GFP+ cINs over the total number of fate-mapped GFP+ cINs) across the different cortical layers of the control and Mef2c cKO (Dlx5/6::Cre;Mef2c^loxp/loxp^RCE) animals. Mef2c results in a reduction in Pvalb density in all cortical layers except for layer 1; Error bars reflect s.e of the mean; P < 0.05*, P < 0.01**, P < 0.001 ***. **H)** Quantification of densities of various cIN subtypes (number of [peptide marker]+ expressing cIN/area) in the P21 control and Mef2c cKO somatosensory cortex. Immunohistochemistry was performed for Pvalb, Sst, Vip, NPY, and calretinin (CR). Error bars reflect s.e of the mean; P < 0.01**

Strikingly, our integrated analysis also revealed strong evidence of interneuron specification in embryonic samples. All four cardinal classes of interneurons exhibited transcriptomically defined precursors by E18.5 (Fig. 4C), revealing novel early embryonic markers for Pvalb, Sst, Id2, and Vip subsets, and also exhibiting transcriptomic evidence of finer subdivisions (Fig. 4E, Supplementary Fig. 14). Moreover, in examining the earliest stages, we observed a separation of Pvalb and Sst-precursor cells within the E13.5 postmitotic population (Fig. 4D), which included a subset of transcriptomic markers that were conserved onward to adulthood (Early marker genes for Pvalb neurons: *Mef2c, Erbb4, Plcxd3*; Early marker genes for Sst neurons: *Sst, Tspan7, Satb1*) (Fig.4E; Supplementary Figure 11). A minority of E13.5 cells also mapped to Vip and Id2 subsets, but conserved transcriptomic markers did not pass statistical significance until E18.5 (E18.5 markers of Vip neurons: *Vip, Synpr, Igf1*; E18.5 markers of Id2 neurons, *Reln, Mpped1, Id2*). Similarly, subtype segregation within the four cardinal classes became evident at different developmental timepoints. For example, the clear emergence of Sst, Id2 and Vip and Id2 subtypes was readily apparent for a subset of cells at E18.5 (Supplementary Figure 13), but we were unable to clearly subdivide the Pvalb subclass at P10, consistent with its late development. Importantly, the results of our integrated analysis (guided by shared sources of variation) was consistent with independent analysis of each developmental dataset (Supplementary Fig. 12), but provided the means to link particular interneuron classes and subtypes to the specific precursor populations that give rise to them. By contrast, we observed no shared sources of heterogeneity indicating the specification of adult subtypes within mitotic progenitors, consistent with our previous results (Fig. 2).

Beyond providing early markers for specific cardinal classes, we wished to determine whether these genes are essential for the emergence of specific interneuron populations. In this regard, the selective expression of *Mef2c* which, based on our findings discriminates early Pvalb-precursors from other MGE-derived interneuron subtypes, represents an attractive target to examine this question. Making this gene of particular interest, genetic variants associated with the transcription factor Mef2c are linked to autism, intellectual disability and schizophrenia^41–43^. We thus sought to validate a functional role for the transcription factor Mef2c in the emergence of the Pvalb cardinal class. Conditional deletion of *Mef2c* in inhibitory neurons using *Dlx5/6::Cre;Mef2c^loxp/loxp^* and *Lhx6i::Cre;Mef2c^loxp/loxp^* led to a specific loss of interneurons by P20 in cortical layers 2-6 (Fig. 4F-H). Thus, not only does it provide an early marker for this cardinal class, *Mef2c* is essential for the generation of this population. Intriguingly, we found that a subset of embryonic cardinal class markers from our dataset (including *Mef2c*) were also differentially expressed in adult human interneurons, based on a recently published single nucleus RNA-seq dataset from postmortem human brain tissue^44^ (Supplementary Fig. 15). A marker list defining embryonic cardinal classes is therefore likely to contain other determinants of interneuron fate determination and maintenance, potentially extending across species.

## DISCUSSION

Our work reveals the genetic trajectories linking GE progenitors to mature subtypes in the adult cortex, demonstrating how subtype specific heterogeneity progresses from the sparse expression of cardinal genes to the emergence of the dazzling array of specific subtypes that populate the mature cortex. As such this work informs us as to the logic underlying cIN diversity. Our data demonstrate that cell cycle and differentiation (i.e. MT) account for the great majority of transcriptional variability within mitotic GE progenitors, while progenitor heterogeneity across eminences is evident in the expression of transcription factors that seed regional identity. Upon becoming postmitotic, three distinct precursor states were observed across all GEs. Cells passing through the first gave rise to interneurons, while cells passing through the remaining two appeared to be destined to generate different projection neuronal subtypes. While branch-specific genes are indicative of a common precursor state, each branch is further characterized by the expression of eminence-specific ‘cardinal genes’ that seed the unique class and subtype identities that arise from the MGE and CGE. It seems probable that the superimposition of precursor state genes and eminence-specific genes cooperate in bestowing both the common and unique characteristics within particular GABAergic populations. Thus, precursor genes likely direct the developmental cascade and acquisition of general properties that are shared within a given class. This likely ensures, for instance, that interneurons migrate tangentially to the cortex or hippocampus, while projection neurons remain positioned ventrally and form long range projections. Supplementing these more general programs are the eminence-specific genes that, for example, may direct the axons of parvalbumin cINs to form perisomal baskets and the efferents of somatostatin cINs to reliably target dendrites. These distinct differentiation modules are reflective of the four major cardinal classes of cortical interneuron precursors. We were able to chronicle the progressive appearance of specific subtypes from these cardinal classes, all of which show clear evidence of transcriptomic separation within the embryo. As subtype differentiation progressively appears within all cardinal classes, the degree to which these cells are covertly committed to a specific subtype within the GEs versus only acquiring such character postnatally when reaching their settling position, remains an open question. Nonetheless, our ability to identify very early precursors that belong to a cardinal class or subtype and the unique genes that define them represents a significant advance. It both offers insight into when specific cell types are generated and provides genetic access to immature cortical interneuron subtypes. Future work is needed to similarly understand the refinement of GE projection neuron subtypes from precursor states. Finally, our findings indicate that components of the transcriptional networks underlying IN fate specification are conserved between mouse and human. This suggests detailed knowledge of the specific cell types that express disease genes, such as *Mef2c* within the early Pvalb cardinal class, will be informative for understanding the etiology of human psychiatric disorders, and points to the exciting potential for cross-species comparisons in both developing and adult neuronal populations.

## ACKNOWLEDGEMENTS

We thank members of the Fishell and Satija Labs, and Claude Desplan, for valuable feedback and discussion, Lana Harshman, Bernadette Bracken, and William Stephenson for assistance with scRNA-seq experiments, and Naomi Habib for assistance with published datasets. This work was supported by NIH P01 NS074972 (GF), R01s NS081297 (GF), MH071679-12 (GF & RS), NIH DP2-HG-009623 (RS), EMBO ALTF 1295-2012 (CM), DFG Postdoctoral Fellow (CH), NIH F30MH114462 (RB), NIH F31NS103398 (KA), and NSF DGE1342536 (AB). GF is also support by a grant from the Simons foundation.

